# The mutation rate in human evolution and demographic inference

**DOI:** 10.1101/061226

**Authors:** Aylwyn Scally

## Abstract

The germline mutation rate has long been a major source of uncertainty in human evolutionary and demographic analyses based on genetic data, but estimates have improved substantially in recent years. I discuss our current knowledge of the mutation rate in humans and the underlying biological factors affecting it, which include generation time, parental age and other developmental and reproductive timescales. There is good evidence for a slowdown in mean mutation rate during great ape evolution, but not for a more recent change within the timescale of human genetic diversity. Hence, pending evidence to the contrary, it is reasonable to use a present-day rate of approximately 0.5 × 10^−9^ bp^−1^ yr^−1^ in all human or hominin demographic analyses.

---

Population genetics provides a theoretical framework for inferring evolution, including changes in demography, based on genetic variation between individuals. It is primarily concerned with relative changes, in the sense that properties such as divergence time and population size are expressed in scaled units whose relationship to the time in years or number of individuals involved is not fixed. This is appropriate for genetic data, which is generally comparative in nature and carries no explicit record of absolute time or population size. However such data is only one of several sources of information about the evolutionary past, and the question of a timescale must be addressed if we want to relate genetic inferences to evidence from fossil, archaeological and paleoenvironmental data.

Most demographic analyses are based on differences due to genomic mutational events, typically single-nucleotide polymorphisms, and the quantities they estimate are naturally expressed as genetic divergence in units of substitutions per base pair. In simple terms, the genetic divergence *d* between two samples can be converted to a time *t* in years since their common ancestor by the expression 2*t* = *d/μ* where *μ* is the mean yearly germline mutation rate over that period. Unfortunately, the question of what value of *μ* to use is less straightforward, as the germline mutation rate depends on multiple factors which may have varied substantially over time, and about which we may have little or no historical information. It also depends on which regions of the genome are analysed and at what level of sensitivity and specificity, making it potentially difficult to estimate an appropriate rate for a given demographic analysis or to compare estimates made using different approaches.

## Mutation rates in present-day and recent human evolution

The first estimates of the human mutation rate predate the availability of molecular genetic data, and were based on the incidence of *de novo* (uninherited) disease cases where the causative allele was thought to be dominant [1, 2]. In recent years, taking advantage of developments in genome sequencing technology, several new methods of estimation using genomic data have been implemented (Figure 1). Of these, estimates of the present-day genome-wide mutation rate have mostly agreed with each other, even as sequencing technologies have developed and sample sizes have grown. In particular, estimates based on whole-genome sequencing in family trios (the majority of studies) have consistently fallen in the range 1.1−1.3 × 10^−8^ bp^−1^ [3–12], as did the first estimate based on identity by descent (IBD) within a pedigree [13]. Other studies have yielded slightly higher estimates however, including a more recent population-IBD estimate which obtained a value of 1.66 × 10^−8^ bp^−1^ [14], and alternative approaches using calibration against different genetic mutational processes [15, 16]. Since sequencing in families detects mutations accumulated over a generation or two at most, whereas other methods are sensitive to somewhat older timescales, one possibility is that the latter reflect higher ancient mutation rates which have slowed in recent human evolution.

However, there are also reasons why sequencing family trios may slightly underestimate the present-day mutation rate. The main advantage of this approach is that potential samples are plentiful, allowing the measurement not just of mean rate but also variation with factors such as parental age and genomic distribution [8, 20]. Also, unlike in other methods the temporal baseline (usually a single generation) is unambiguous. Its principal disadvantage is that single-generation *de novo* mutations are rare relative to the error rate in variant calling (60-100 mutations per individual), so false negative and false positive rates are both high and difficult to estimate. To mitigate this, genomic regions where variants are difficult to call are generally excluded via filtering, but these regions are not easy to identify and the callable genome length may be overestimated, leading to an underestimate in the per-bp mutation rate. Most of the studies cited here have attempted to quantify and account for this using simulations or validation against other methods of variant discovery, but it remains possible that true *de novo* mutation rates are consistently underestimated to some degree.

Another potential downward bias arises from the fact that trio sequencing experiments generally compare somatic cells rather than germ cells. An early post-zygotic mutation occurring prior to germline specification in either parent may be detected in that individual’s soma as well as his or her offspring, and hence, seemingly present in both generations, might not be correctly identified as a *de novo* mutation [21–23]. This could be a significant factor if cellular mutation rates are particularly high in the earliest cell divisions of embryogenesis.

**Figure 1:**
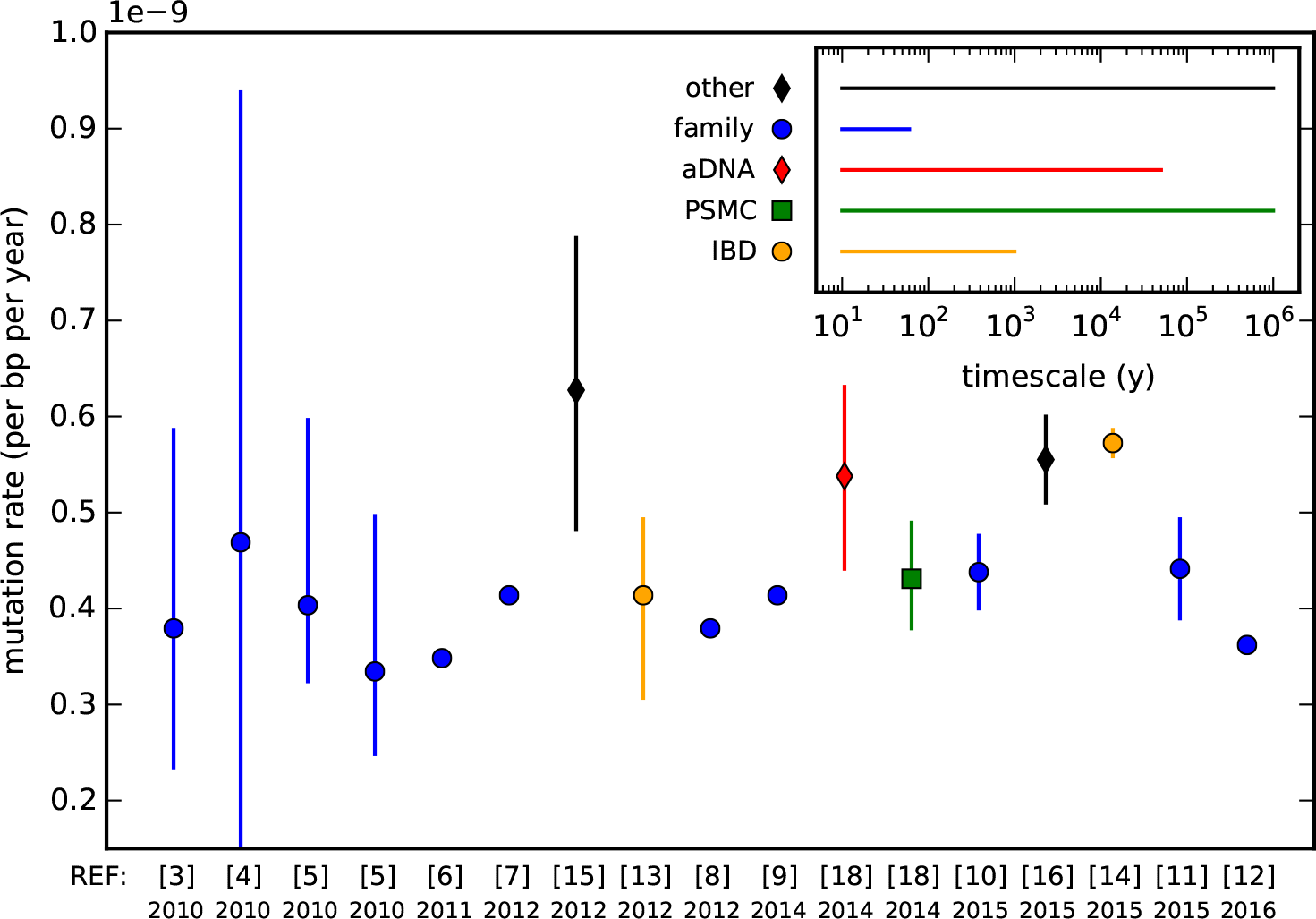
Recent estimates of the human genome-wide mutation rate. Estimates are shown as yearly rates, scaled where necessary using a mean generation time of 29 yrs [17]; confidence intervals (90% or 95%) are shown where reported. Citation numbers and publication years are given on the *x*-axis. **family**: Family sequencing compares genomes sampled from consecutive generations in one or more families, and within each one identifies *de novo* mutations present in offspring and in neither parent [3–12]. Per-generation mutation rate is calculated as the mean number of de novo mutations seen divided by the length of ‘callable’ genome sequenced (the number of genomic positions where a de novo mutation would have been called if present). **IBD**: Estimation based on identity by descent (IBD) detects de novo mutations as differences between chromosomal tracts which have been inherited IBD within or between individuals, for example in samples which are related to each other within a multi-generation pedigree. Information about the number of generations separating chromosomes may come from genealogical records [13] and/or from genetic inference [14]. **aDNA**: Estimation based on branch shortening in ancient DNA uses genome sequence data from an ancient human sample of known age (established with radioisotope dating) and divides the mean number of extra mutations found in present-day humans by the separation in time [18]. **PSMC**: The pairwise sequential Markovian coalescent method infers ancestral effective population size from diploid genome sequence data [19]. A mutation rate can be estimated as the one which best aligns effective population size histories inferred from modern and ancient samples after accounting for the known age difference between them [18]. **other**: Methods based on comparison with other mutational clocks: calibration using coalescent time estimates based on microsatellite mutations [15]; calibration against the recombination rate and expected variation of heterozygosity in diploid genomes [16]. *Inset:* Indicative timescales over which mutations detected by each method (or which otherwise influence its estimate) have accumulated.

In principle, estimates based on IBD in a multi-generation pedigree should be less susceptible to either of these biases. Multiple accumulated mutations in IBD tracts are more easily distinguished from sequencing noise than in family sequencing, especially for larger pedigrees, and this approach can detect all germline mutations (excepting perhaps early post-zygotic mutations in the common ancestor of a given tract). However they are not without their own methodological issues: genealogical information and uncertainty in the inference of relatedness and IBD are potential sources of error. In particular the boundaries of IBD tracts and the path of inheritance may be ambiguous, and the total extent of regions in which mutations can be detected may be quite limited except in close and inbred pedigrees. Pedigree datasets are also more difficult to collect, and since the two such genome-wide estimates published to date have not overlapped [13, 14], it is difficult to assess the significance of their disagreement with other methods. We can expect forthcoming studies to help clarify this picture.

### The exomic mutation rate

Mutation rates are known to vary between genomic loci [24], and the estimates discussed above are based either on whole genome data or (in the case of IBD estimates) on regions sampled genome wide without regard to location or context. Other studies, mostly using family trio sequencing, have been based on data sampled only from exomes [25–30], and have tended to yield higher values than equivalent whole-genome studies (ranging from 1.3-2.0 × 10^−8^ bp^−1^ with a mean of 1.5 × 10^−8^ bp^−1^). This is consistent with the elevated GC content of genic regions and the increased mutability of GC-rich sequence (discussed below), but it may also be that the biases discussed above are of less consequence for exome sequencing data. Consistent with the latter possibility is the fact that a recent IBD-based estimate in exomes of 1.45 × 10^−8^ bp^−1^ [31] is only slightly below the mean of trio-based estimates published so far.

## Mutation rates in great ape evolution

Before the advent of high-throughput genome sequence data, estimates of the human mutation rate were generally based on phylogenetic calibration: *μ* = *d/2t_s_*, where *d* is the genetic divergence between two species and *t_s_* the time since speciation as estimated from the fossil record. In principle, allowance must also be made for the difference between speciation and genetic divergence times, corresponding to coalescence within the ancestral population, but in practice the magnitude of this can usually only be guessed [32]. Phylogenetic calibration has some potential advantages: fossils can often be dated with relatively high accuracy using radiometric or stratigraphic methods, and since it estimates the mean substitution rate over the time separating the two species, it accounts automatically for selection and other time-varying factors which may complicate extrapolation from present-day rates.

By the time the first mutation rate estimates from *de novo* sequencing appeared, the field had largely settled on a consensus value of 1.0 × 10^−9^ bp^−1^ yr^−1^ for the yearly rate in hominid evolution [33]. Thus the finding that *de novo* estimates were a factor of two lower than this prompted considerable debate [34]. For some events, such as the speciation of humans and chimpanzees, a higher rate had been increasingly difficult to reconcile with fossil and archaeological data, and a lower value (implying older date estimates) mostly improved concordance [35, 36]. However for more ancient events a longer timescale was problematic, and to a large extent remains so still. For example, applying a present-day human mutation rate of 0.5 × 10^−9^ bp^−1^ yr^−1^ to the 2.6% genetic divergence between humans and orang-utans [37] yields an divergence time of 26 Mya. Even allowing for a large ancestral coalescent time of 5 Myr this is substantially older than the dates of 12-16 Mya typically quoted in paleoanthropological literature [38]. The difference increases for older dates: the human-macaque divergence [39] implies a speciation more than 40 Mya, whereas paleoanthropological studies generally place this node at 25–30 Mya [40].

One way to resolve this discrepancy is to regard it as evidence for a faster mutation rate 20 Mya or more, and hence a slowdown in mean rates since that time. In fact, such a hypothesis is also supported by differing branch lengths within the primates as measured from an outgroup or common ancestor, with hominid (great ape) lineages being shorter than those of other primate groups by a factor of 1.4–1.6 [41–43]. Because the branch shortening applies to all great apes (albeit to varying degrees), such a slowdown cannot have occurred only on the human branch, and if it occurred more recently than the hominoid ancestor (so that a higher mean mutation rate applies to dating the orang-utan divergence), it must have involved a degree of parallelism across all hominid lineages. This is not impossible for closely-related species, but might be regarded as unlikely *a priori*. It is estimated that compared to humans, chimpanzees have evolved only 2% faster since divergence, and gorillas 7% faster [41]. Indeed, a measurement of present-day mutation rate in chimpanzees based on sequencing *de novo* mutations in a multigeneration pedigree has also produced a value of 1.2 × 10^−8^ bp^−1^ [44], very similar to equivalent human estimates.

However, timing constraints based on fossil evidence should be handled with caution. Even where fossils are themselves well dated, their correct placing relative to a particular speciation event may be far from straightforward [45, 46]. There may also be important differences between the evolution of anatomical phenotypes represented in fossil taxa and the genetic differences involved in speciation, particularly when the possibility of ancestral population substructure around the time of speciation is taken into account. More fundamentally, fossil evidence tends to be more informative about lower bounds than upper bounds on speciation dates (essentially because the presence of derived characteristics is more informative than their absence), and so ‘stem’ taxa which appear ancestral to a speciation event provide only weak constraints on its earliest possible date [45, 47, 48]. Thus it may be premature to conclude that a genetic estimate of 20–23 Mya for the orang-utan speciation is irreconcilable with fossil evidence, and the implied slowdown in mutation rate may be less than expected both in magnitude and (especially if prior to the orang-utan speciation) in the degree of any parallel evolution involved.

## Causes and correlates of mutation rate variability

In addition to direct evidence from present-day and ancient genomic data, it may also be possible to learn about past mutation rates indirectly by studying the underlying physiological and population genetic factors affecting them. Some understanding of these factors, and how they may have varied in the past, comes from considering the cellular origins of germline mutation.

Germline mutations can arise from disruption of the DNA molecule at any time within a germline cell, but most are believed to result from errors in DNA replication during cell division, referred to as replicative mutation [49]. Over multiple generations the rate of replicative mutation will depend strongly on the mean number of cell divisions from zygote to zygote, and this can differ between species, between sexes, and perhaps also between populations due to variation in reproductive behaviour. Differences between the germline in males and females reside primarily in the sex-specific nature of gametogenesis [50], where there is a much greater number of cell divisions on the paternal lineage due to the fact that spermatogenesis involves a continuous process of stem cell division throughout adult life. This in turn contributes to a greater accumulation of *de novo* replicative mutations passed on by the father than by the mother, a phenomenon referred to as the male mutation bias and found in many species, with important evolutionary consequences [51]. Its effect in humans has been quantified in recent sequencing studies, with estimates of the male/female ratio in mean number of transmitted mutations ranging from 3.1-3.9 [8, 9, 12, 15].

A further consequence is that the older the father, the more cell divisions his gametes will have passed through, and hence the more mutations they are likely to carry. The resulting age effect in paternally transmitted mutations has been measured at 1.2-2.0 additional *de novo* mutations per year of paternal age in recent studies [8–10, 15, 52], corresponding to a doubling from puberty to age 30. In fact this is substantially less than expected under the standard model for spermatogenesis [50, 53], which predicts a factor of ten increase over the same period based on the number of cell divisions involved. Possible explanations for this discrepancy include a revised model of spermatogenesis in which gonial stem cells pass through fewer cell divisions, or strong variation in per-cell-division mutation rates during development, with much higher rates prior to gametogenesis [11, 23, 53].

Sequencing studies initially measured no significant age effect in maternally transmitted mutations [8, 9, 15], consistent with replicative mutation under the longstanding reproductive model in which, after proliferation during fetal development, oocytes are held in stasis until maturation later in life and experience no postnatal cell divisions [54]. However, two more recent studies have estimated significant effects amounting to 0.51–0.86 additional mutations per year of maternal age [12, 52]. The initial negative findings may have resulted from methodological factors (for example, strong correlation in the data between maternal and paternal ages makes the effect difficult to discover if information on parent of origin for *de novo* mutations is not available). The distribution of parental ages sampled, which can can differ even between large cohorts, particularly at the extremes of the distribution, has also been suggested as an important factor [12, 53, 55].

The measured maternal age effect is weaker than the paternal effect, but nevertheless supports the view that aspects of our longstanding model of gametogenesis need revision. In fact, evidence for a maternal age effect in larger-scale mutations such as chromosomal abnormalities has been available for some time [56, 57], motivating the ‘production line’ hypothesis for oogenesis [58], which attributes the effect to a correlation between the number of pre-natal cell divisions experienced by oocytes and the age at which they are matured for ovulation. Other hypotheses have been proposed however [59], including the possibility that previously undetected post-natal oogenetic cell divisions may occur, analogous to the gametogenetic process in males [60, 61]. This is supported by the discovery of germline stem cells in the ovaries of adult female humans and mice [62], and thus perhaps also by the finding of a non-zero maternal age effect in genomic mutations.

Alternatively, or in addition, non-replicative or spontaneous germline mutations may play a greater role than is generally assumed and contribute to age effects in both sexes [63]. Such mutations can arise from instability or disruption of the DNA molecule itself, for example due to oxidative mutagens within the nucleus or exposure to ionising radiation. Unlike replicative mutation, we might expect spontaneous mutations to accumulate on germline lineages at a rate which is independent of the number of cell divisions or life history parameters such as generation time, and hence to behave more like a molecular clock. (Purely clock-like behaviour is perhaps unlikely however, as the production of oxidative mutagens is a causative factor for spontaneous mutation and is itself proportional to metabolic rate, which also scales with generation time [64].) It has been shown that the relative contribution of spontaneous mutation depends to a large extent on the efficiency of DNA repair [49]: if such repair is rapid relative to the length of the cell cycle then most spontaneous mutations will be corrected prior to replication, and replicative processes will dominate. This is believed to be the case for most mutation on the human germline; however this is largely due to the correspondence between the paternal age effect and the number of mitotic divisions in males, an assumption which is perhaps undermined by the finding of a non-negligible maternal effect (assuming female post-natal mitoses are negligible).

There are also genomic loci where spontaneous mutation is expected to play a dominant role, notably CpG sites, in which the cytosine when methylated (as is usually the case in mammals [65]) is prone to spontaneous deamination from C to T. (As an aside, if spontaneous mutation contributes substantially to the maternal age effect we might expect an even stronger effect at CpG sites [49]. This was not observed in the only study so far to have examined it [12], but larger studies in future may have greater power to detect a difference.) The more clock-like behaviour of CpG mutations is borne out in branch length comparisons within the primates and other mammals (for example, root-to-tip distances vary by 2-4 times less than for other mutational types) [41, 42, 66]. This makes CpG sites potentially appealing for ancestral demographic inference. However they are rare in the genome, particularly in intergenic regions (1% of sites genome-wide and 3% in exons, where they have presumably been maintained by purifying selection) [67], so their use in this way is limited to site-wise analyses ignoring haplotype information. Moreover their behaviour is not strictly clock-like but only more so than other mutations, so branch-specific factors must still be taken into account.

## Discussion

The first proposed solution to the mutation rate problem was the molecular clock hypothesis of Zuckerkandl and Pauling [68], essentially a zeroth-order approximation which ignored rate variation, yet which proved surprisingly successful (in Crick’s words, ‘much truer than people thought at the time’ [69]). In the decades since, the quest to improve upon this approximation has focused primarily on calibration against the fossil record, using increasingly sophisticated models to account for rate variation and stochasticity in fossil creation and discovery [70]. Notwithstanding the advances made in this direction, it is clear that the recent accessibility and availability of genome sequence data in humans and other species has opened a new window on the germline mutation rate.

It is also clear that generation time alone, while important, is insufficient to fully describe the dependence of mutation rates on developmental and reproductive processes. Germline mutation depends on a plurality of related biological timescales: the ages of puberty and reproduction, the duration of fertility and of key stages in embryonic development, the cycle times of cellular processes in gametogenesis, and the efficiency of DNA repair, each potentially differing by sex or species [23, 53, 71, 72]. The sequencing studies discussed here have begun to explore these phenomena, and although some initial findings have differed or disagreed, further insights into their present-day effects and how they might have varied in the past can be expected from future sequencing on population scales. Important evidence for ancestral reproductive behaviour and life history parameters may also come from paleontological and archaeological data [73–75], and more direct evidence continues to come from ancient DNA. In particular, a recent analysis has shown that the mean generation time has not changed appreciably over at least the last 45,000 years, based on the rate of decline in linkage disequilbrium resulting from Neanderthal admixture in several ancient human samples [76].

We return therefore to the question of what mutation rate to use in analyses of human demographic evolution. Figure 1 provides a weak indication that methods sensitive to older mutation events tend to yield higher estimates, but this is somewhat confounded with potential downward bias in whole-genome estimates from family sequencing. Branch length comparisons within the apes provide no support for a substantial human-specific slowdown [41]. It may be that future developments will reveal recent modest changes in mutation rate, perhaps differing between modern human populations [31, 77], driven by evolution in one or more of the factors discussed here, and possibly more substantial differences in other hominins if data becomes available. Pending such refinements however, a reasonable (and conservative) approach is to apply a yearly mutation rate of 0.5 × 10^−9^ bp^−1^ yr^−1^ uniformly to analyses of demographic events within or between human populations, including between modern and archaic humans.

Finally, and notwithstanding that there are many gaps in our understanding, it is worth noting that the role of the mutation rate in human demographic inference has changed markedly in recent years. Whereas genetic data were formerly regarded as definitive about topological relationships between taxa but uninformative about their timescale, this distinction has vanished or even reversed in the case of recent human evolution. Estimates of the mutation rate have begun to converge, and it has become clear that many events in human demographic history are more complex than previously assumed, with populations diverging gradually or in convoluted ways with ongoing gene flow or later admixture [78–81]. It is fair to say that in many, perhaps even most cases, the mutation rate is no longer the principal source of ambiguity in human demographic inference.

## Acknowledgements

I am grateful for support from an Isaac Newton Trust/Wellcome Trust ISSF Joint Research Grant.

